# Remote Learning Barriers and Opportunities for Graduate Student and Postdoctoral Learners in Career and Professional Skill Development: A Case Study

**DOI:** 10.1101/2020.10.07.328492

**Authors:** Margery Evans Gardner, Elizabeth C Bodiya, Shoba Subramanian

**Author notes:** Denotes corresponding author Contact information.

## Abstract

Career and professional development competencies are critical for biomedical PhD and postdoctoral training. In the current educational landscape, programs that meet these competencies are offered and attended in an ad hoc manner. During the COVID-19 pandemic and the accompanying switch to virtual learning, our team observed a surge in interest for our weekly non-sequential programs. In this study, we surveyed our learners to better understand motivators for attending these programs during the pandemic and to identify barriers for participating in such events before and during Work-From-Home. Our data indicate that conflict with research responsibilities, time spent to get to the event location, and planning time to attend are all significant barriers to engagement. Notably, feelings of being overwhelmed, which increased slightly during the pandemic, stood out as an identified barrier. Per our results, the virtual format was an attraction. While 58% of respondents would prefer to access professional development programs virtually in the future, almost 42% indicated a preference for in-person events when normalcy resumes as the physical presence of an instructor and of peers result in a deeper engagement. Our collective analysis here suggests that learners will benefit from a hybrid or combination of synchronous and asynchronous career and professional development programming in the future, even post-pandemic, to reduce identified barriers. Alongside hybrid learning engagements, we strongly recommend structured time for learners to enhance their professional competencies, enabled by a commitment from departments and faculty mentors to enable equity in professional skill building and fostering lifelong growth mindset.

## INTRODUCTION

Our team supports the <u>Career and Professional Development (CPD</u>) of the ∼1200 graduate and postdoctoral learners at the University of Michigan (UM) Medical School. We provide a variety of programs ranging from weekly workshops and seminars to structured, time-limited, and cohort-based programs. These evidence-based programs, which follow published recommendations (1–4), are aimed at driving professional, academic, and career success in the biomedical sciences and cover a gamut of topics including career exploration, transferable skill building, job application best practices, and portfolio building. While our programs are extra-curricular and not required to meet degree requirements, participating learners find significant value in them as reported in our <u>2019 Annual Highlights</u>.

Most universities across the USA shut down in March 2020 due to the rapidly increasing COVID-19 cases. Leadership units at research-intensive universities focused on two main items: a) pivoting credit-bearing curricular courses to online formats with some structural support (example <u>communication</u> from UM President for Spring and Summer courses going remote) and b) developing and implementing procedures for prioritizing critical research activities and enabling a rapid research ramp-down (5). Beyond the early years of PhD training, there is seldom any requirement for course-based learning for PhD and postdoctoral learners. Instead, biomedical and life sciences PhD students and postdoctoral fellows spend most of their working hours in their training laboratories conducting inquiry-based research. Thus, the research ramp-down, in combination with the lack of structured learning requirements, created a lot of undefined and unplanned time and space for these learner populations. While faculty advisors used their discretion to provide some structure to learners’ research through remote data analysis and writing manuscripts and grant proposals, this likely varied between labs and advisors.

In our state of Michigan, the <u>“Stay Safe, Stay Home”</u> mandate alongside that from UM in mid-March began a period of time hereafter referred to as Work-From-Home (WFH). Until WFH, our offerings had been exclusively in-person. We rapidly reimagined our programs to enable remote learning. Through virtual programs, we continued to meet our learners’ needs. Additionally, we created new programs covering many aspects of written and visual science communication to meet the likely demand for these skills during WFH and to guide learners’ unstructured time in a purposeful way. Peer units on our campus and similar teams across the country also pivoted CPD programs from in-person to remote formats.

Upon launching our remote program offerings, we observed a dramatic increase in RSVP numbers as compared to when our programs were offered exclusively in-person. In order to understand motivators for remote attendance to CPD programs and to pinpoint specific barriers in program participation before and during WFH, we conducted a systematic survey. In this manuscript, we share key findings of this survey. These findings, and our evidence-based recommendations that stem from them, inform future practices for CPD programming. This includes formats and content for UM and peer institutions across the country. We believe that these recommendations, which can easily be implemented with limited budgets, enable more flexibility, inclusion, and equity for diverse learners to access critical programs, albeit extra-curricular, more effectively and efficiently.

## METHODS

### CPD Marketing and Communication

All events were marketed via email using the Mailchimp marketing platform. UM Medical School graduate students and postdoctoral fellows are added to the distribution list during onboarding. Additional subscribers outside the Medical School are allowed to opt in. The distribution list consists of approximately 800 graduate students, 1,200 postdocs, and 500 faculty and staff University-wide. The total number of recipients varies as appointments change or as opt-in or out updates happen. Non-medical school-affiliated are included in these numbers. Individual event announcements were sent approximately one week prior to the event. Events were also marketed in a CPD weekly newsletter and listed on our website. Learners could RSVP for events through the individual event announcements, newsletter, or the website.

### RSVP and Attendance Data Collection and Analysis

RSVPs were collected digitally using either Google Forms or Sessions at Michigan (an internal event management tool developed by the UM Office of the Vice President for Student Life). During registration, learners were asked to indicate their stage of training, program, and are asked to submit a question or topic they wish to see addressed during the given event. During earlier CPD offerings, attendance was collected manually using a paper sign-in sheet. The Sessions at Michigan platform allows for digital attendance collection for which an Apple iPad was used. Attendance to virtual events during WFH was collected using the Sessions at Michigan Self-Check-In feature using a shortened URL or QR code. Staff members also cross-referenced the virtual attendee list with the Sessions at Michigan registration to log attendance. All registration and attendance data in this study were collected using the method described above with one event facilitated by an outside vendor as the only exception. Statistical significance was established comparing before and during WFH RSVP and attendance rates each using a 2-tailed unpaired Student’s t-test.

### Survey Design and Analysis

The survey created for this study is under IRB exemption (#HUM00187729) from UM Office of Research. The survey was created using the UM Qualtrics XM survey platform. An electronic link to the survey with informed consent was distributed to PhD and Master’s students, post-baccalaureate scholars, and postdocs (approximately 1,200) within the medical school using an internal listserv. The anonymous survey was active for 8-9 days. The total number of surveys collected was 249 representing approximately 20% of the learner population. The survey contained a variety of questions related to how learners engaged in CPD activities before and during the WFH order from the State of Michigan (enacted March 17, 2020). Questions also probed factors that motivated learners to participate or served as barriers to participation in CPD activities before WFH and during WFH. Motivators to participation were scored on a four-point scale: not a factor, somewhat of a factor, strong factor, or N/A (not applicable). The data presented in Figure 3 represent all “somewhat of a factor” and “strong factor” responses. Respondents selected whether each barrier to participation was a factor before WFH, during WFH, or neither. Respondents could indicate that a barrier was present both before and during WFH. To assist in future planning, learners were asked to indicate if they would prefer to engage in in-person or virtual activities and what their motivations were for either. The survey concluded with several optional demographic questions: stage of training, years in current position, gender identity, underrepresented minority, and nationality.

**Figure 1:**
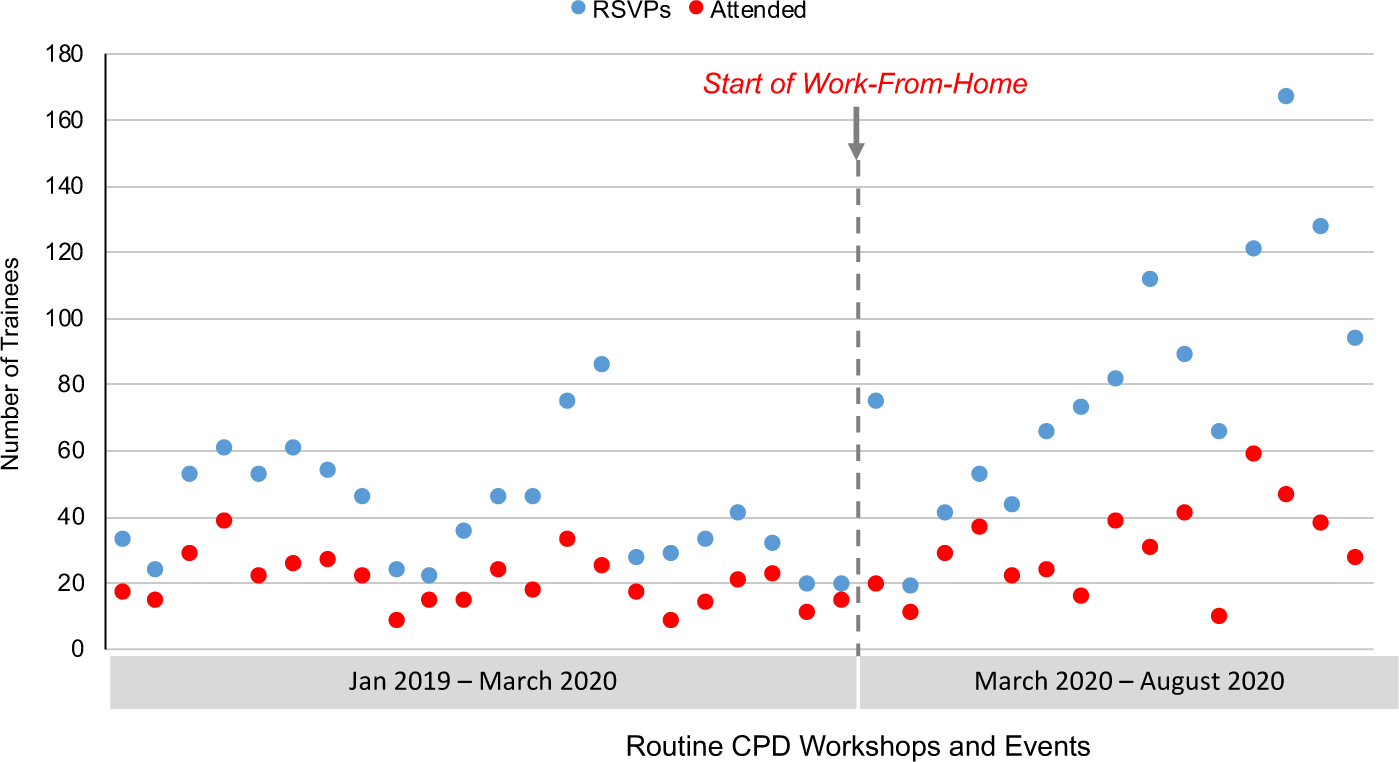
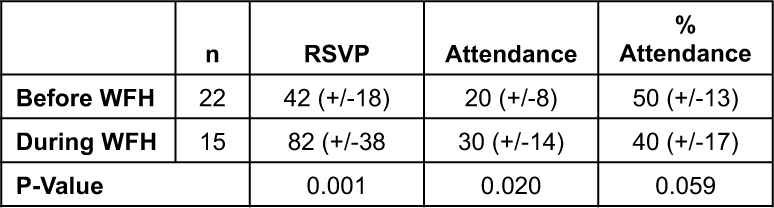
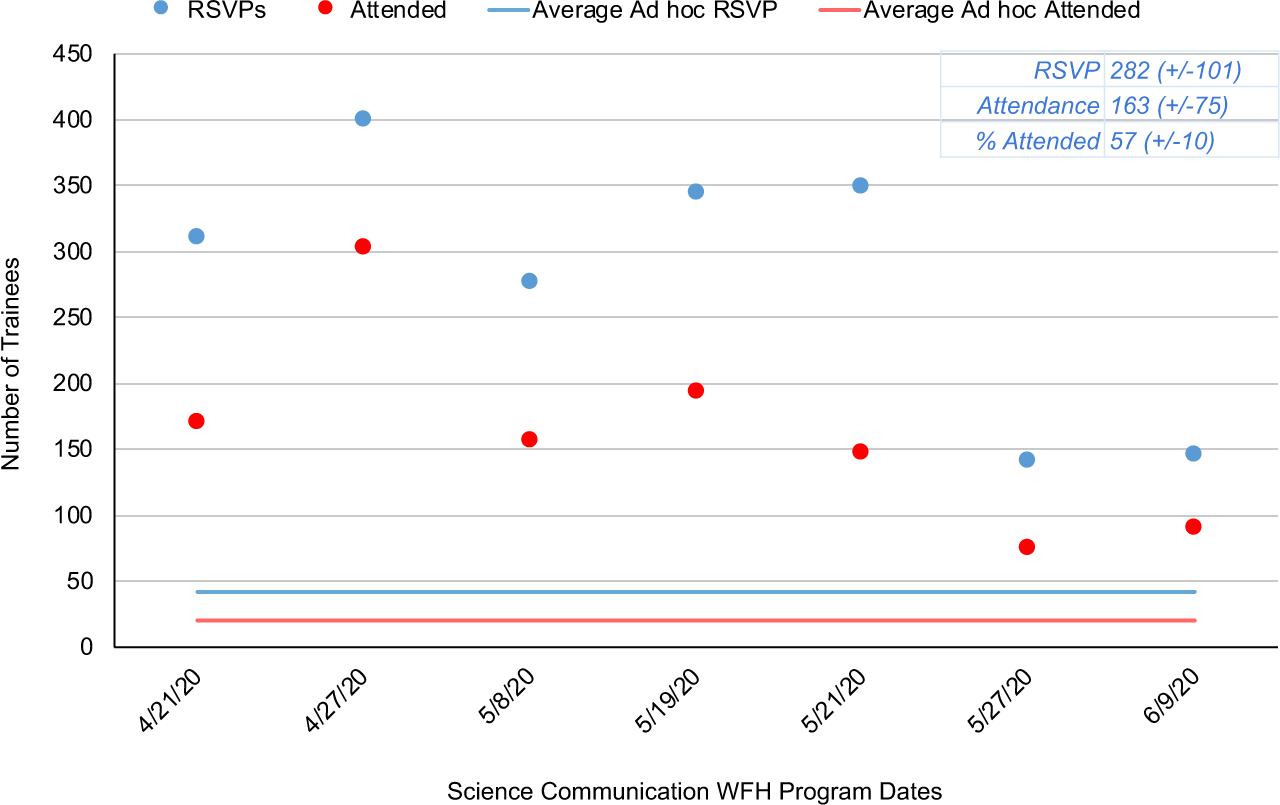
Interest and Attendance Increased in Remote CPD Programming. A. Learner RSVPs (blue) and attendance (red) for routine, ad-hoc CPD. Transition from in-person to virtual programming at the start of Work-From-Home is indicated by the grey arrow and dotted line. B. Program RSVPs and actual attendance numbers were collected for all in-person and virtual programs before and during WFH. Shown are averaged +/− standard deviation. P-values were calculated using a 2-tailed unpaired t-test. C. Learner RSVPs (blue circles) and actual attendance (red circles) for science communication programming during WFH is compared to the average RSVP rate for regular, ad hoc events (blue line) and actual attendance (red line). Average RSVPs, actual attendance, and % of RSVPs who attended are reported in the inset.

**Figure 2:**
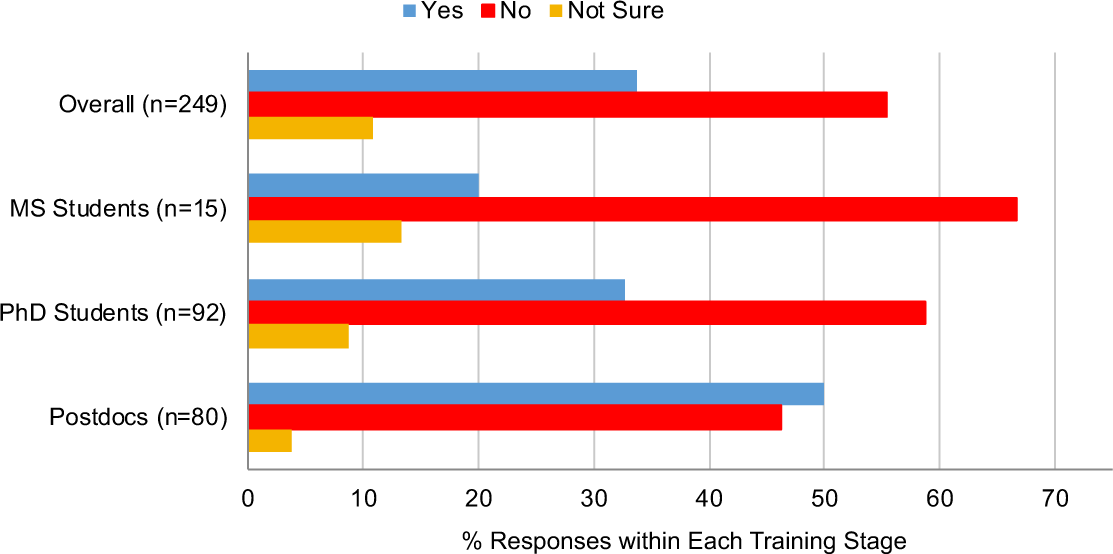
Learner Perceived Frequency of Attendance in CPD Programming During Remote Learning and WFH. Responses divided by learner stage and presented as a percentage of the total responses within the respective training stage. Learners reported their perceived attendance frequency in CPD programs during in-person learning and virtual WFH learning. Learners responded “Yes” to increased participation in CPD programming (blue), “no” their participation did not increase (red) or if they were “not sure” (yellow).

**Figure 3:**
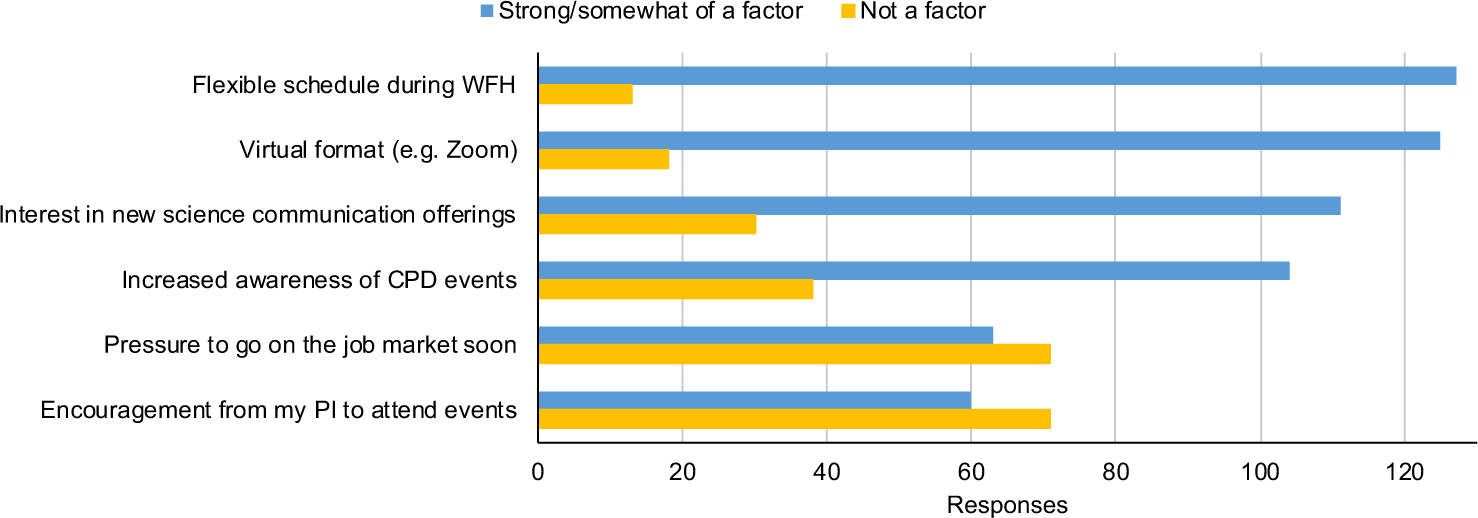
Primary Motivators for Increased Participation in CPD Programming During Remote Learning and WFH. Learner responses (n=145, x-axis) on the level of influence by each of six motivators (y-axis) on participation are shown. Respondents could indicate whether each motivator was “strong factor” in participation, “somewhat of a factor”, “not a factor”, or “not applicable”. Respondents who selected motivators as a “strong factor” or “somewhat of a factor” in participation were pooled. Respondents who identified motivators as “not applicable” are not shown.

## RESULTS

At the Office of Graduate and Postdoctoral Studies, we have been offering weekly programs for career and professional development (CPD) tailored for our graduate and postdoctoral learners since January 2018 (<u>refer to our events calendar</u>). Since the WFH mandate was announced at UM, we carefully pivoted all our programs to remote formats primarily using video conferencing software such as BlueJeans and Zoom. During this mandate, we observed changes in RSVP and attendance numbers for similar weekly, non-cohort-based programs when compared to before WFH (Figure 1A). There was an increase in RSVP with an average 42 RSVPs/event before WFH and 82 RSVPs/event during WFH (Figure 1B). Interestingly, although the average percentage of RSVPs who actually attended the event went down from slightly (51% pre- to 40% during-WFH) it was not a statistically significant decrease (Figure 1B). Additionally, we piloted a couple of science communication workshops at the beginning of remote learning, which gained RSVPs of 300-400/event. Based on this significant interest, we conducted a needs assessment check-in to understand this considerable uptick. We identified manuscript writing, peer review, and scientific speaking as further topics of interest. As a result of these, we observed an average of 282 RSVPs/event with a 57% attendance rate across all science communication topics offered (Figure 1C).

Based on the increased interest in participation from mid-March – August 2020, we were curious to a) identify the primary motivators that encouraged increased participation and b) better understand the barriers that kept graduate and postdoctoral learners from attending CPD events before and during WFH. To that end, we created a short survey that addressed the above questions along with other queries on preferred formats of programming in the future, even after the pandemic/WFH restrictions have been lifted. The survey was sent out via email to our entire learning community of Master’s, PhD, and postdocs at the UM Medical School.

First, we asked whether the frequency of the respondent’s engagement in CPD programs increased during WFH (Figure 2). 50% of postdocs indicated that they had increased their frequency of CPD participation during WFH whereas about 4% were unsure. In the case of Masters and PhD students, fewer respondents said that they had an increase in the frequency of attending CPD programs during WFH compared to those that did not. Overall, however, 34% of our respondents indicated that they attended more professional development events during WFH and remote learning.

Next, we wanted to understand what the motivators were for learners to attend remote events during WFH. This question was filtered to include only respondents who answered “yes” or “not sure” to whether their participation increased during WFH. As seen in Figure 3, approximately 125 respondents selected either virtual format or schedule flexibility during WFH as a strong or somewhat of a factor for attendance. Not surprisingly, interest in science communication events obtained the third highest selections, which is corroborated by the initial needs-assessment survey for new programs during WFH. This was followed by increased awareness of events as a motivator. Pressure to go on the job market and encouragement from PI (Principal Investigator/research advisor) were ranked below these other factors, but still received a significant number of responses.

Beyond the motivators for learner participation in remote events, we sought to understand the barriers that impacted learner participation before and during WFH. Across nine barriers listed in the survey, with an additional ‘other’ category, learners were asked if each barrier impacted their ability to participate in CPD activities before WFH, during WFH, or neither. Respondents could select both before and during WFH. Before WFH, transportation/location of events, conflict with date/time, and conflict with research responsibilities were significant barriers to participation, with the top barrier impacting close to half of the respondents (Figure 4A). Our data show a reduction in each of these barriers during WFH. On the other hand, feeling overwhelmed more than doubled, caring for dependents went up five-fold, and lack of stable internet went up sixteen-fold as barriers during WFH. Personal conflicts, awareness of programs, and PI support to attend events remained mostly similar before and during WFH.

**Figure 4:**
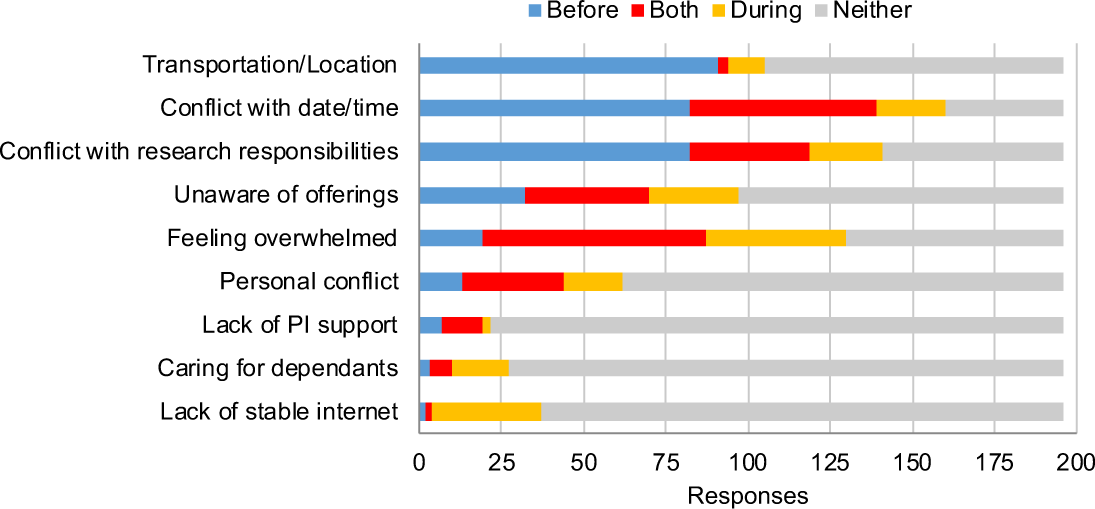
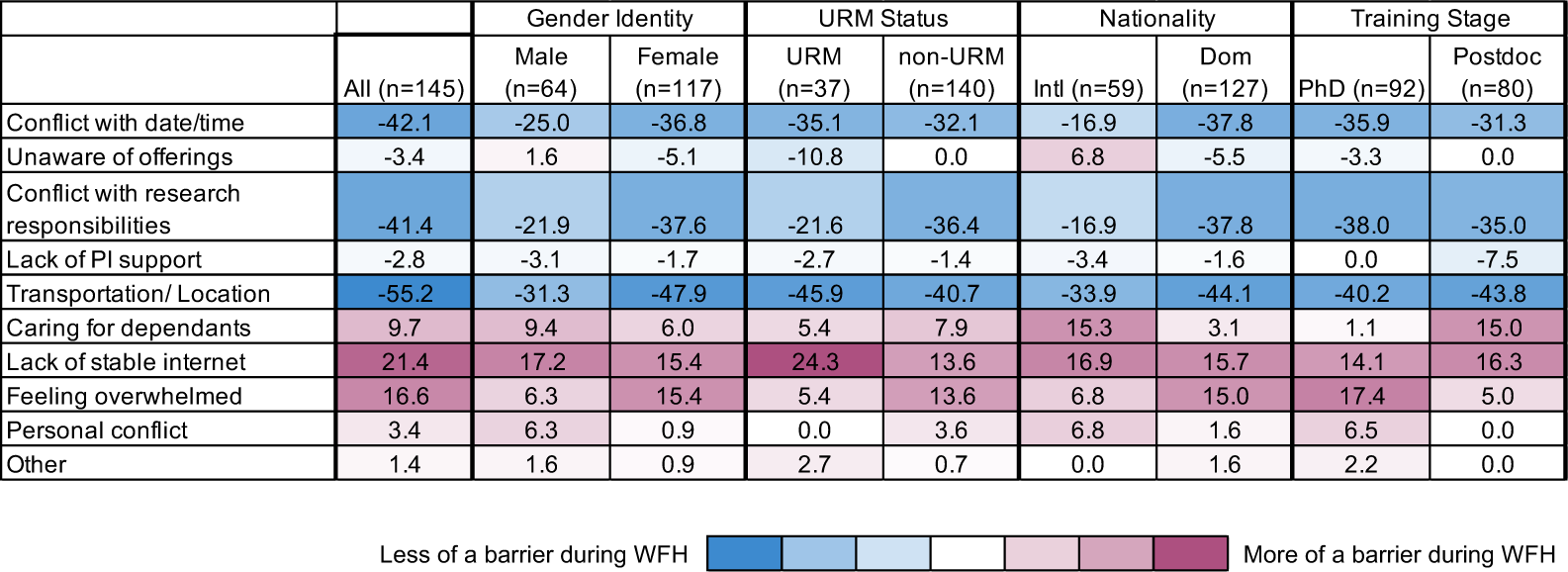
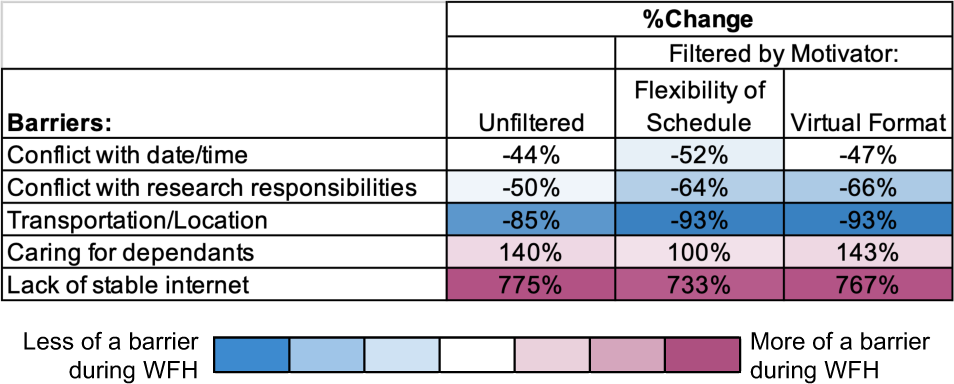
Primary Barriers for Participation in CPD Programming Before and During WFH. A. Learner responses to whether nine barriers (y-axis) were a factor in their ability to participate in CPD activities before WFH (blue), during WFH (yellow), both before and during WFH (red) or not a factor during either time period (grey). Total responses for each selection are reported (y-axis). B. Barriers encountered by learners were broken down by demographic or self-described identity. Responses in this table include barriers before WFH, during WFH, and both. Values reported are the change in percent of responses within each demographic normalized to the total number of responses within the demographic (n). Demographic reporting was voluntary. Data shown only for respondents who self-identified. (URM = Under-Represented Minority; Intl = International; Dom = Domestic; PI = Principal Investigator). Data from learners identifying as non-binary gender identifies were not included as the number of responses was too small to maintain anonymity. C. Responses to three barriers that decreased during WFH and two barriers that increased during WFH were filtered to only include those who indicated that increased flexibility of schedule or virtual format during WFH was a motivator to attending virtual CPD programming.

Next, we compared changes based on several self-identified demographics in the barriers for CPD event attendance before and during WFH, including those who chose some barriers for both periods (Figure 4B). While the trends within each demographic largely reflected the overall trends, there were some variations between groups. The number of female respondents who reported feeling overwhelmed during WFH increased by 15.4% compared to their male counterparts who only increased by 6.3%. Barriers related to caring for dependents disproportionately affected postdocs and international learners during WFH. Also, a greater number of learners who identified as an Under-Represented Minority (URM) reported that the lack of stable internet was more of a barrier during WFH compared to non-URM learners. Fewer male respondents reported that barriers due to research responsibilities were alleviated by WFH compared to female respondents. Interestingly, fewer URM (21.5%) and international (16.9%) learners reported that barriers related to their research responsibilities were relieved during WFH compared to their non-URM (36.4%) and domestic (37.8%) counterparts, respectively.

Based on our findings that several barriers to participate in CPD programming differ before and during WFH, we asked whether there is a correlation between the top barriers before WFH and the top motivator of schedule flexibility for attending virtual events during WFH. As seen in Figure 4C, the top motivating factors (flexibility of schedule and virtual format during WFH) correlated very strongly with a decrease in the top three barriers that are identified between before WFH and during WFH (including respondents who chose both time periods): date and time conflict, research responsibilities, and transportation to the event location.

Besides understanding the recent motivations and barriers in CPD participation, we also asked our learners about their preferences for the format of future programming. Our survey asked learners if they would prefer to mostly engage in remote or in-person events, when in-person events are a safe option once again. Across learner groups, 58% of respondents indicated a preference for remote events, while 42% selected a preference for a return to in-person programs, as highlighted in Figure 5A. Trends per learner group closely followed the same preference. Respondents who selected remote/virtual programming for the future (n=108) were asked for their top three reasons driving this preference. Figure 5B shows that 77% noted that they would not lose time on cross-campus transportation by attending virtual programming. Other top reasons included the ability to watch recorded sessions afterward (62%) and that remote options helped learners prioritize CPD (58%). Almost half of respondents, 48%, indicated that they learn equally as well or better in remote situations. Of those who would prefer future remote programming, 31% indicated a higher level of comfortability when engaging virtually. Lower factors in virtual preference included learners utilize inclusive teaching features (e.g. closed captioning) in remote learning (19%) and learners who juggle caregiving duties (8%). Write-in answers further emphasized location and distance, with some learners living outside of Ann Arbor, thus remote options would be more accessible to them.

**Figure 5:**
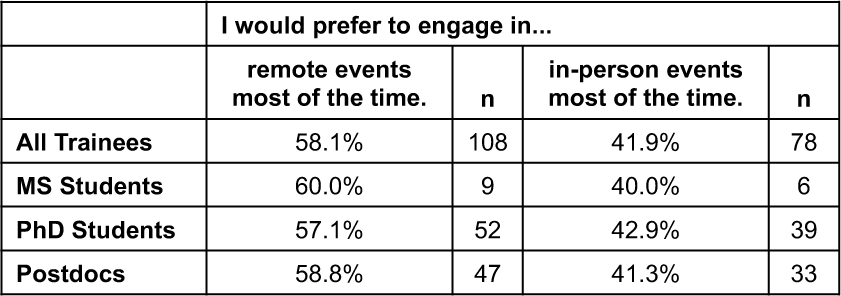
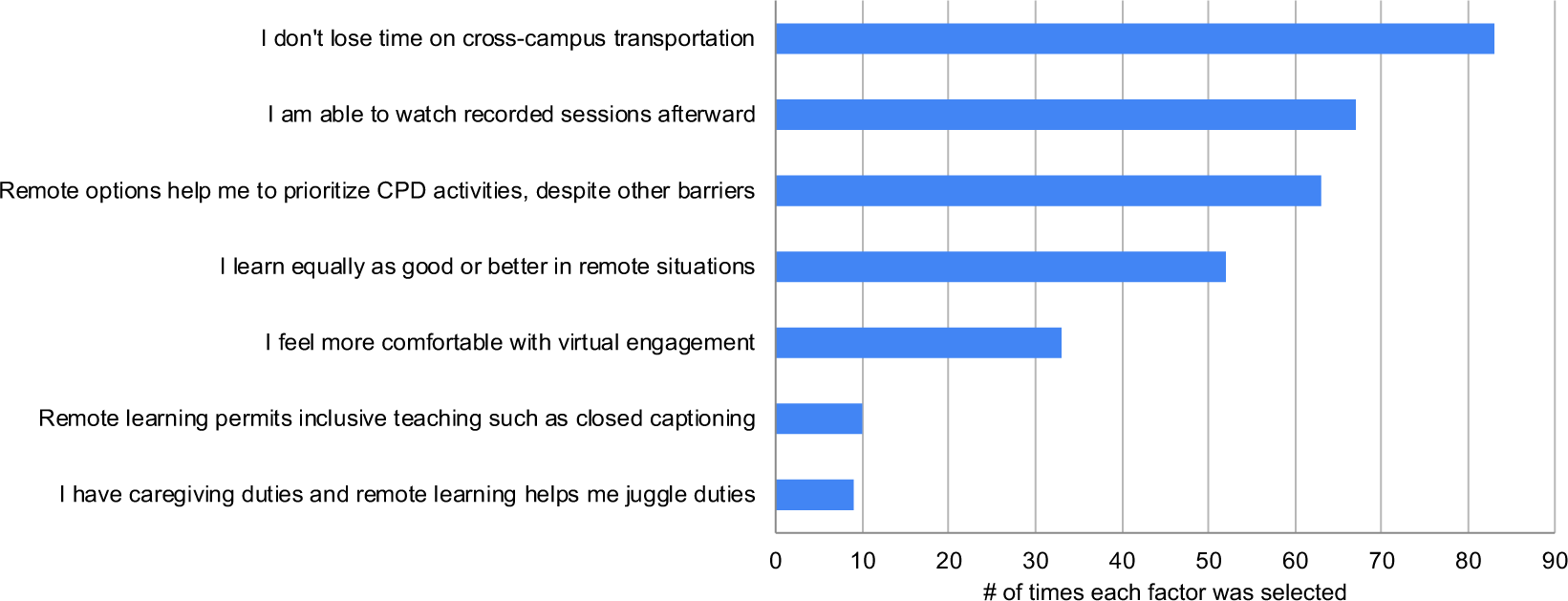
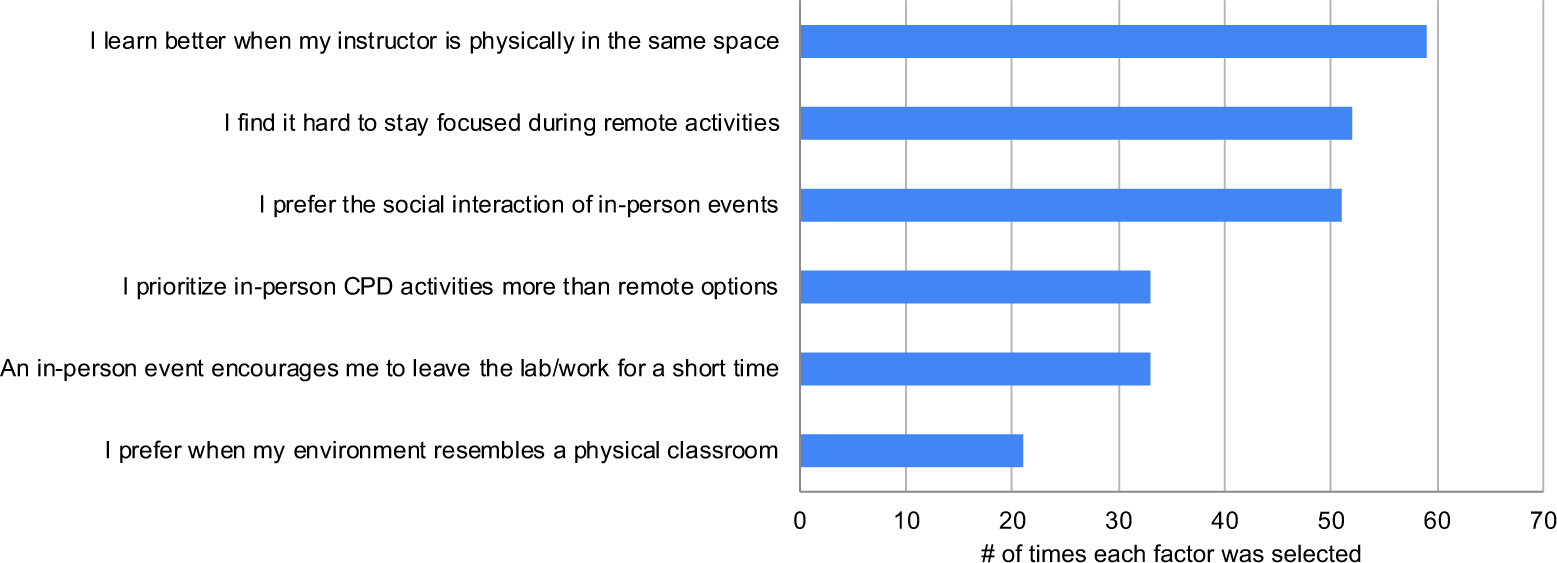
Learner Preference and Reasoning for Preferred Format of Future CPD Programming. A. Learners responses (n=186) on preference for future CPD program format in either remote or in-person events. Responses are separated by training stage. B. Learners who prefer future remote CPD programming (n=108) chose the top three reasons among seven choices (y-axis) for their preference. Reported are the number of times each reason was selected (x-axis). C. Learners who prefer future in-person CPD programming (n=78) chose the top three reasons among six choices (y-axis) for their preference. Reported are the number of times each reason was selected (x-axis).

Figure 5C displays why 40% of learners prefer a return to in-person CPD. 76% indicated that the physical presence of an instructor improved learning. Other top reasons referenced the challenge of staying focused when attending a remote session (67%) and a desire for social interaction (65%). Over 40% of those preferring in-person sessions shared that they prioritize in-person activities more than remote options, and in conjunction, in-person programs encourage the physical requirement to leave the lab and work for a short time (42%). To a lesser degree, 27% of those that favor future in-person events prefer an environment that resembles a physical classroom, including the presence of peers.

## DISCUSSION

Career and professional development (CPD) training of graduate and postdoctoral learners is pivotal in the changing landscape of career outcomes (6, 7). Targeted transferable skill building and active career planning is key for future workforce development (8). Significant strides have been made across several institutions and federal funding agencies to prioritize quality and quantity of CPD programs (9). These programs are typically offered by several offices ranging from graduate schools, college-level graduate and postdoctoral training offices, as well as departmental units. Despite the advances, most training programs do not require CPD program participation toward meeting any required milestones in education. This in turn meant that when the pandemic-driven shut down of research and associated WFH mandate was put in place, it was left up to the individual units how they wanted to continue with their career and professional development programming.

Our team regularly communicates with learners to understand needs and accordingly adapts CPD programming to meet them. Therefore, tailoring programs at the beginning of remote learning was not an unprecedented move on our part as it was primarily based on learner feedback. However, the significant spike in RSVP for our routine events and science communication workshops was rather surprising. Based on this participation increase and learner feedback, it was clear that they found high value in these programs. We further believe that they used our space to find purpose during a time when everything else was mostly undefined.

Interestingly, although there was an increase in participation per event during remote learning, fewer than 50% of the survey respondents said that they increased their frequency of participation. This can be explained either by the possibility that a) we attracted many of the same learners who in the past attended our program or b) it alludes to a bias in who responded to the survey or c) it points at a likely misalignment in learner perception. Such a misalignment can be attributed to the fact that learners may not have the necessary calibration to compare a before/after timepoint, since CPD is not curricular and is accessed in an ad hoc manner.

From the responses we gathered, it was unequivocal that major barriers exist for learners in planning these events into their daily schedules. These barriers are exacerbated on large, multi-campus institutions like ours, where labs are spread out as far as 3 miles away, and transportation/parking are limited, which could be amplified in the winter months. Interestingly, almost the same number of respondents selected transportation/location as a barrier before WFH as those who indicated it was not a barrier during either time. This suggests that learners from labs proximal to routine program venues are able to attend more easily than others. While the location issue could theoretically be solved by offering programs in multiple locations, in many campuses access to reserving classroom spaces to meet capacity and other requirements are often a huge premium. Additionally, some event spaces require an internal fee. Learner access to parking continues to be a huge limitation, making it equally difficult for learners who are able to use their own vehicles to commute to the events. Finally, like many of our peer institutions, our CPD programming is supported by a small staff with limited capacity to run events in multiple locations. These barriers collectively drive systemic inequities in learners’ ability to access in-person programs.

Conflict with research responsibilities was indicated as a major barrier to attending CPD events before and, to a lesser extent, during WFH. This conflict, at some level, is at odds with national recommendations from the National Academies of Science, Engineering, and Medicine (3, 10), the Council for Graduate Schools (3), and the National Postdoc Association’s <u>Core Competency List</u>, all of which strongly suggest that CPD programs be integral to learning in order to build skills for developing future workforce. While those learners on federal training grants make targeted plans for CPD requirements and to meet funding expectations, the intentional planning or participation mandates are not the same for others. Thus, without changes in programmatic elements and infrastructural support, the barriers to balance research with CPD program engagement will continue to limit the holistic development of graduate and postdoctoral learners. This is of particular importance as several barriers disproportionately impact learners in marginalized demographic groups. While international learners reported conflicts with date, time, and research responsibilities decreased during WFH, this decrease was less than half what domestic learners reported. This could be because domestic learners are better aware of research expectations and balancing time. This brings up the idea of a “hidden curriculum” wherein learners new to (American) Higher Education systems are not aware of expectations that are not overtly communicated (11). This also disproportionately affects first-generation learners (12). The inequities revealed between URM/international learners and non-URM/domestic learners around barriers due to research responsibilities suggests that international and URM learners felt or were more compelled to continue research activities despite research ramp down and the WFH mandate. The barriers faced included access to critical WFH resources such as stable internet, which impacted more URM learners during WFH than non-URM learners, and assistance in caring for dependents, which affected more postdocs and international learners. Many of these inequities could be ameliorated by creating resources including protected time aligned with clear expectations for competency development within the training experience for all learners to engage in professional development activities not limited to coursework and research. An intentional professional development curriculum will not only drive success in a learner’s future career but will also enhance academic success and outcomes in their training years.

Science communication training including the core areas of written communication including manuscript writing, creating figures, peer review best practices occurs within the research labs with ad hoc guidance by PIs or senior lab members. The unprecedented interest and attendance we had for these events indicates that the historical apprenticeship-based training where communication skill building and feedback is primarily dependent on advisors or labs is not sufficient. Our data indicate that this is an area of immediate and high need that should be formalized into the training journeys of both graduate and postdoctoral learners.

As we plan for a future when returning fully to in-person work is not a barrier, we find that the split between learners who prefer remote learning as compared to in-person is small. Importantly, the reasons picked for either preference align with their personal motivations and barriers. Moreover, these data shed light on other systemic issues. For example, postdoctoral fellows place higher weight on the physical instructor presence and on peer social interaction when they indicate an in-person preference. This aligns with postdoctoral training years being isolating for many learners (13, 14).

In the future, we recommend institutional infrastructures for graduate and postdoctoral CPD learning to have core features for flexibility, accessibility, and inclusion, as shown in Figure 6, to complement other published recommendations (15). Ultimately, CPD can be easily adapted for remote learning compared to many other aspects of biomedical graduate and postdoctoral training. Despite this, it has not been at the forefront of learning goals for advanced STEM learners. Herein, based on our experiences delivering remote programs during a pandemic, we have uncovered new barriers as well as corroborated previously known issues. Based on learner voices, we use this opportunity to provide institutional recommendations toward making CPD more accessible to all learners.

**Figure 6:**
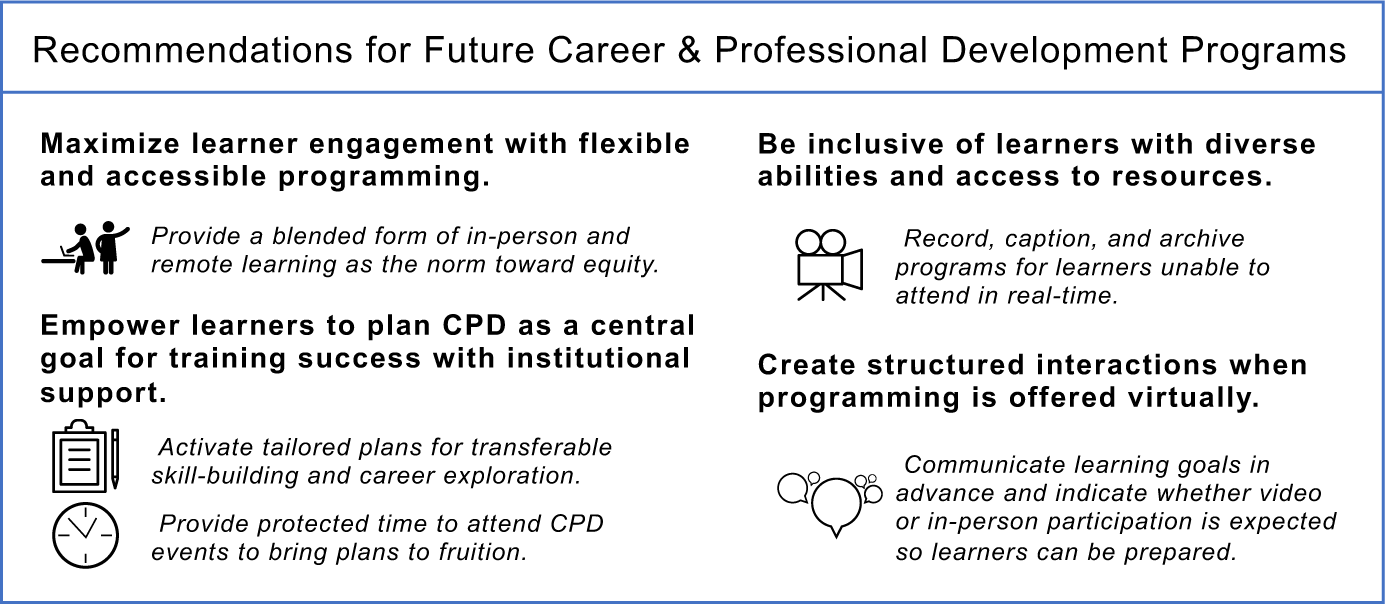
Future Recommendations: To serve as guidelines for practitioners, administrators, and institutions toward enabling flexible, inclusive, equitable, and engaging Career and Professional Development (CPD) programs for graduate and postdoctoral learners.

Finally, the barrier of “feeling overwhelmed” was unambiguous in learners reporting this feeling before and during WFH. Such feelings among graduate students have been discussed in another recent study after the pandemic (16). Although this feeling can easily be attributed to the overall mood of 2020, which has been beset with multiple calamities converging at once, learners need dedicated and proactive support in mental health and well-being at every stage of their training and under all circumstances (17), so they can show up to engage fully. Our recommendations to maximize CPD planning into the daily lives of busy learners, if turned into action, may at least alleviate anxieties around career prospects and build confidence in our future leaders.

## ACKNOWLEDGEMENTS

The authors thank Dr. Manoj Puthenveedu and Dr. Kaylee Steen for critical feedback on the manuscript. We also thank past and present team members at the UM Medical School’s Office of Graduate and Postdoctoral Studies under the leadership of Dr. Mary O’Riordan and appreciate all of our campus partners at UM. This work is partly funded by an NSF IGE grant (#1954967) awarded to Dr. Subramanian.

